# Many bat species are not potential hosts of SARS-CoV and SARS-CoV-2: Evidence from ACE2 receptor usage

**DOI:** 10.1101/2020.09.08.284737

**Authors:** Huan Yan, Hengwu Jiao, Qianyun Liu, Zhen Zhang, Xin Wang, Ming Guo, Bing-Jun Wang, Ke Lan, Yu Chen, Huabin Zhao

**Affiliations:** State Key Laboratory of Virology, Modern Virology Research Center, College of Life Sciences, Wuhan University, Wuhan, 430072, China; Department of Ecology, Tibetan Centre for Ecology and Conservation at WHU-TU, Hubei Key Laboratory of Cell Homeostasis, College of Life Sciences, Wuhan University, Wuhan 430072, China; Frontier Science Center for Immunology and Metabolism, Wuhan University, Wuhan, 430072, China

**Author notes:** Correspondence to: Huabin Zhao,; Yu Chen,; Ke Lan. These authors contributed equally to this work (Huan Yan, Hengwu Jiao).

## Abstract

Bats are the suggested natural hosts for severe acute respiratory syndrome coronavirus (SARS-CoV) and SARS-CoV-2, the latter of which caused the coronavirus disease 2019 (COVID-19) pandemic. The interaction of viral Spike proteins with their host receptor angiotensin-converting enzyme 2 (ACE2) is a critical determinant of potential hosts and cross-species transmission. Here we use virus-host receptor binding and infection assays to show that ACE2 orthologs from 24, 21, and 16 of 46 phylogenetically diverse bat species – including those in close and distant contact with humans – do not support entry of SARS-CoV, SARS-CoV-2, and both of these coronaviruses, respectively. Furthermore, we used genetic and functional analyses to identify genetic changes in bat ACE2 receptors associated with viral entry restrictions. Our study demonstrates that many – if not most – bat species are not potential hosts of SARS-CoV and SARS-CoV-2, and provides important insights into pandemic control and wildlife conservation.

## Introduction

The unprecedented pandemic of COVID-19, caused by the novel coronavirus SARS-CoV-2, has led to major threats to public health and economic development. It is therefore critically important to identify natural or intermediate hosts of SARS-CoV-2 to prevent further spread of COVID-19 and future emergence of similar diseases. Inferred from sequence similarity of human and bat virus genomes, it was suggested that horseshoe bats (*Rhinolophus* spp.) might be natural hosts of SARS-CoV and SARS-CoV-2 (*1-3*). These suggestions have resulted in misguided fears on all bats, and unwarranted attacks on many bats – including species other than *Rhinolophus* – thereby seriously impacting efforts towards bat conservation (*4*). Given the remarkable diversity of bats, which includes more than 1400 species across the globe (*5*), assessing the possibility that diverse bat species act as potential hosts of SARS-CoV and SARS-CoV-2 is urgent and crucial for both controlling outbreaks and protecting populations of wildlife.

ACE2 is the host cell receptor of SARS-CoV and SARS-CoV-2, and plays a vital role in mediating viral entry to cause infection (*1, 6*). The interaction of a virus with its host receptor has been repeatedly demonstrated to serve as a primary determinant of host range (*7*). Here we test ACE2 orthologs from 46 bat species across the phylogeny, including species occurring in urban and in rural areas, for their ability to support the entry of SARS-CoV and SARS-CoV-2. Hence, this study assesses whether diverse bat species are potential hosts of SARS-CoV or SARS-CoV-2. Moreover, by determining the correlation between proximity to humans and probability of being natural hosts of the two viruses, these results provide important insights into pandemic control and wildlife conservation.

## Results

### Evolution of ACE2 in bats inhabiting urban or rural areas

We collected ACE2 orthologs from 46 bat species across the phylogeny **(Figure 1 and Table S1)**. These species contained 28 species that roost or forage in urban areas in close proximity to humans, and 18 species more restricted to rural areas and hence likely to have minimal contact with humans **(Table S2)**. In total, the examined species represent 11 bat families that contain 1345 species, accounting for 96% of all bat species **(Table S3)**. After aligning the protein sequences of bat ACE2 orthologs, we examined 25 critical residues involved in the binding of the surface spike glycoprotein (S protein) of SARS-CoV-2 **(Figure S1)** (*8*). Genetic variations were observed in nearly all these 25 sites, which may have led to different abilities to support entry of SARS-CoV and SARS-CoV-2 (*8*). Furthermore, we detected at least 22 amino acid sites that are putatively under positive selection **(Table S4)**, indicative of heterogeneous selection pressure across sites. Notably, four of these positively selected sites are located in the binding region of ACE2 to the SARS-CoV-2 S protein **(Table S4)**.

**Fig. 1.**
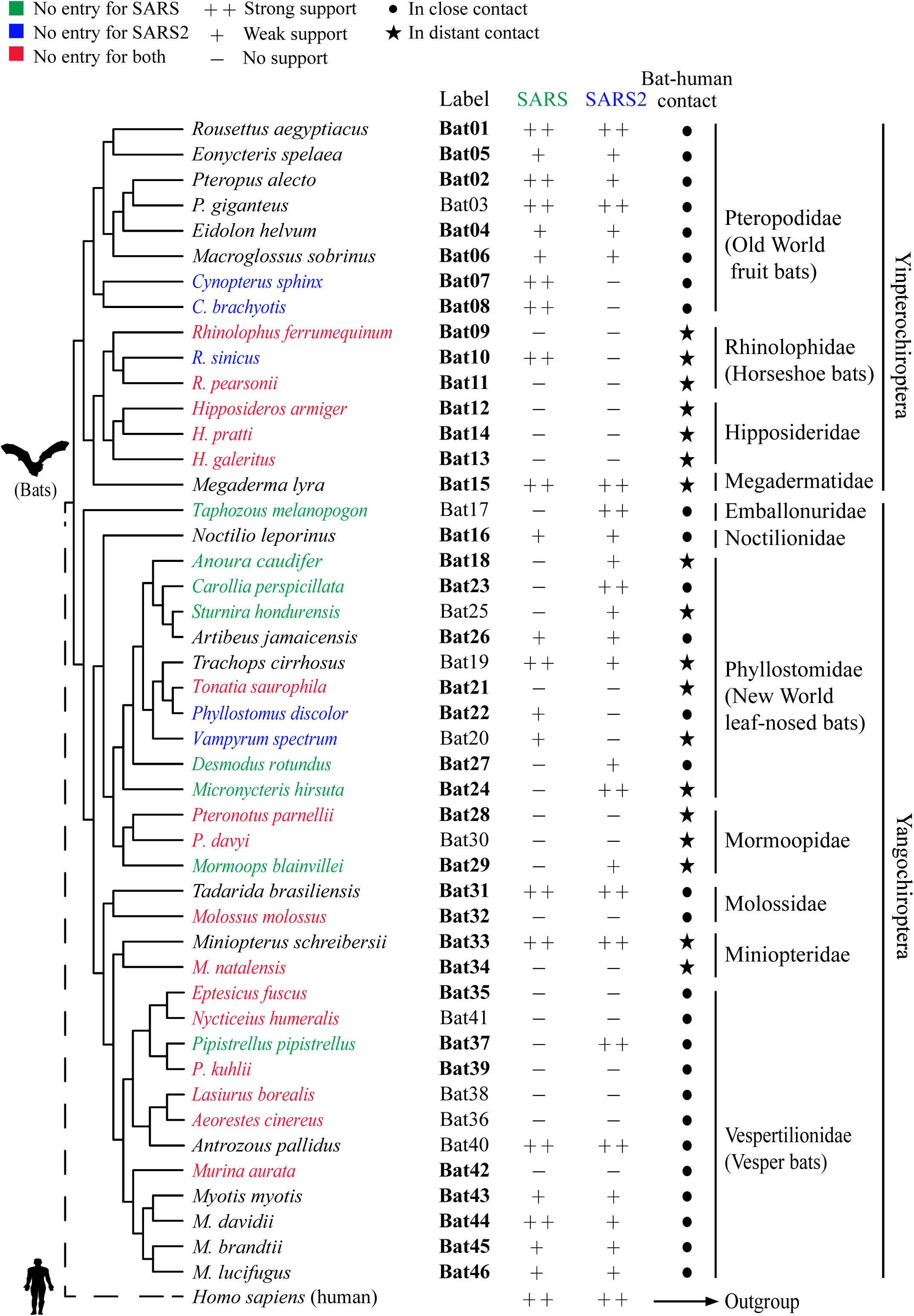
Phylogenetic tree of 46 bat species in this study. Labels of bat species in our experiments are indicated. Infection abilities of bat ACE2 to support SARS-CoV and SARS-CoV-2 entry are shown with different signs: infection data are indicated as % mean values of bat ACE2 supporting infection compared with the infection supported by human ACE2; infection efficiency smaller than 5% is indicated with a minus sign (-), between 5% and 50% a plus sign (+), and greater than 50% a double plus sign (++). Labels shown in bold indicate the bat species that have been examined by in silico analyses in a recent study (*8*). Bat phylogeny was taken from previous studies (*25-27*).

### Interaction between bat ACE2 orthologs and SARS-CoV or SARS-CoV-2 receptor binding domain (RBD)

Efficient binding between the S protein and the ACE2 receptor is important for SARS-CoV and SARS-CoV-2 entry. This binding is mainly mediated by the interaction between the critical residues on the RBD and ACE2. To characterize the receptor function of ACE2 orthologs in a range of diverse bat species, we generated a stable cell library consisting of cell lines expressing the respective 46 bat ACE2 orthologs through lentiviral transduction of 293T cells lacking ACE2 expression (*9*). All bat ACE2 orthologs were exogenously expressed at a comparable level after puromycin selection, as indicated by Western-blot and immunofluorescence assays detecting the C-terminal 3×Flag tag **(Figure 2A-B)**.

**Fig. 2.**
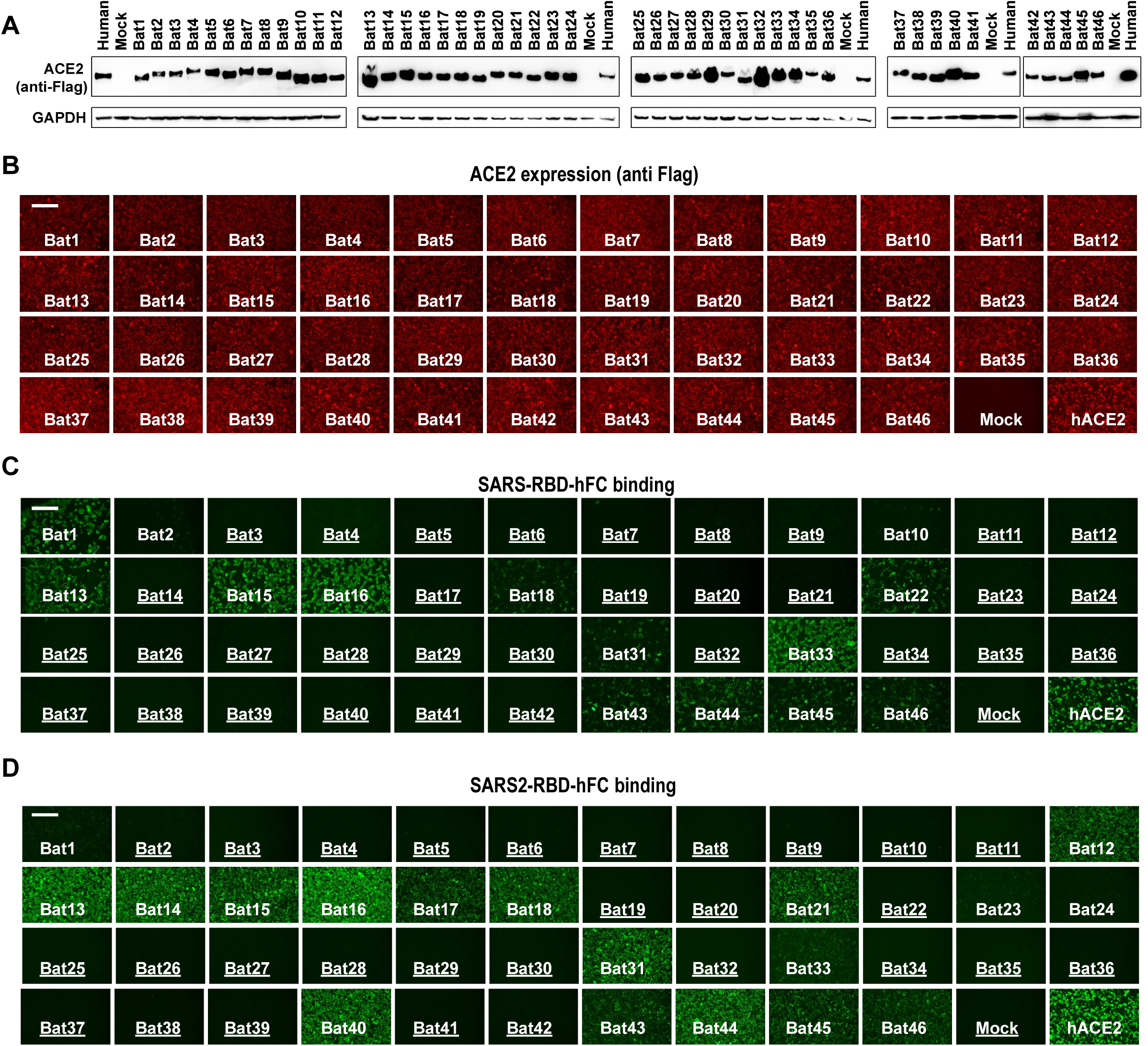
Expression of bat ACE2 orthologs and their interaction with SARS-CoV and SARS-CoV-2 RBD. (A) Western blot detecting the expression levels of ACE2 orthologs on 293T stable cells by targeting the C-terminal Flag tag. Glyceraldehyde 3-phosphate dehydrogenase (GAPDH) was employed as a loading control. (B) Visualization of the intracellular bat ACE2 expression level by immunofluorescence assay detecting the C-terminal Flag tag. Scale bar=100 μm. (C-D) Assessment of the interaction between different ACE2 orthologs and SARS-CoV-RBD-hFc (C) or SARS-CoV-2-RBD-hFc (D). Species that do not support efficient binding are underlined. 293T cells stably expressing the different bat ACE2 orthologs were incubated with 5 μg/ml of the recombinant proteins at 37°C for 1 h. The binding efficiency was examined by Alexa Fluor-488 Goat anti-human IgG through fluorescence assay. Scale bar=200 μm.

To analyze the interaction, we produced recombinant SARS or SARS-CoV-2 RBD human IgG Fc fusion proteins (RBD-hFC), previously reported to be sufficient to bind human ACE2 efficiently (*10, 11*). The protein binding efficiency was tested on the bat ACE2 cell library through immunofluorescence or flow cytometry targeting the human Fc. As expected, binding was almost undetectable on mock 293T cells, but a strong binding signal was detected on the 293T cells expressing human ACE2 **(Figure 2C-D)**. Consistent with previous reports (*12, 13*), SARS-CoV-2 RBD showed higher binding to hACE2 than SARS-CoV, which can also be observed on many bat ACE2 orthologs **(Figure 2C-D)**. Previous reports have shown that only a small fraction of ACE2 orthologs from tested mammalian species could not bind with SARS-CoV-2 S protein [n=6 of 49 species (*7*); n=5 of 17 species (*14*)]. However, our study revealed that many bat species (n=32 and n=28 of 46 species) do not support efficient binding with SARS-CoV-RBD and SARS-CoV-2-RBD, respectively **(Figure 2C-D)**. The overall profiles of bat ACE2 to bind to SARS-CoV and SARS-CoV-2 RBD are generally comparable; a few showed contrasting modes of binding preferences **(Figure 2C-D)**. For instance, Bat22 can bind to SARS-CoV but not SARS-CoV-2, whereas Bat14, 21, 40 can bind to SARS-CoV-2 but not SARS-CoV **(Figure 2C-D)**. Flow cytometry analysis showed consistent results **(Figure S2)**.

Overall, the RBD-hFc binding assays demonstrated that bat ACE2 orthologs showed different affinity and selectivity levels to SARS-CoV and SARS-CoV-2, indicating that ACE2 receptors of many bat species may not support efficient SARS-CoV and SARS-CoV-2 infection.

### Receptor function of bat ACE2 orthologs to support the entry of SARS-CoV and SARS-CoV-2 using pseudotyped and live viruses

To further evaluate the receptor function of different bat ACE2 orthologs, we employed a Vesicular Stomatitis Virus (VSV)-based Rhabdoviral pseudotyping system for mimicking the coronavirus spike-protein mediated single-round entry (*14*). SARS-CoV and SARS-CoV-2 pseudotypes were generated by assembling the coronavirus spike proteins and the replication-deficient VSV with the VSV glycoprotein (VSVG) gene replaced with a fluorescence protein (VSV-dG-GFP) or a Firefly Luciferase (VSV-dG-Luc) reporter (*14*). Both viruses showed minimal background infection on 293T cells, but efficient infection on 293T-hACE2 cells **(Figure S3)**. The susceptibility of the 293T cells expressing bat ACE2 orthologs was then examined with SARS-CoV and SARS-CoV-2 pseudotypes. The results showed that bat ACE2 orthologs have varying abilities to support coronavirus entry, and different preferences for SARS-CoV and SARS-CoV-2. **(Figure 3A-B, Table S5)**. Pseudotypes with GFP reporter showed similar results **(Figure S4)**. Notably, we found that 24, 21, and 16 of the 46 bat species showed almost no entry for SARS-CoV, SARS-CoV-2, and both of these viruses, respectively **(Figures 1 and 3A-B, Table S5)**, suggesting that these species are not likely to be potential hosts of either or both of these coronaviruses. The bat species showing no viral entry include those that occur in urban areas as well as those more restricted to rural areas **(Figure 1, Table S1)**, suggesting that there is no correlation between proximity to humans and probability of being natural hosts of SARS-CoV or SARS-CoV-2. Although horseshoe bats were suggested to be potential natural hosts of SARS-CoV and SARS-CoV-2 (*1-3*), only one of the three examined species (*Rhinolophus sinicus*) supported SARS-CoV entry; this species was suggested to be the potential host of SARS-CoV (*3, 15*). None of these tested horseshoe bats showed entry for SARS-CoV-2 **(Figures 1 and 3)**. These results unambiguously indicate that ACE2 receptor usage is species-dependent.

**Fig. 3.**
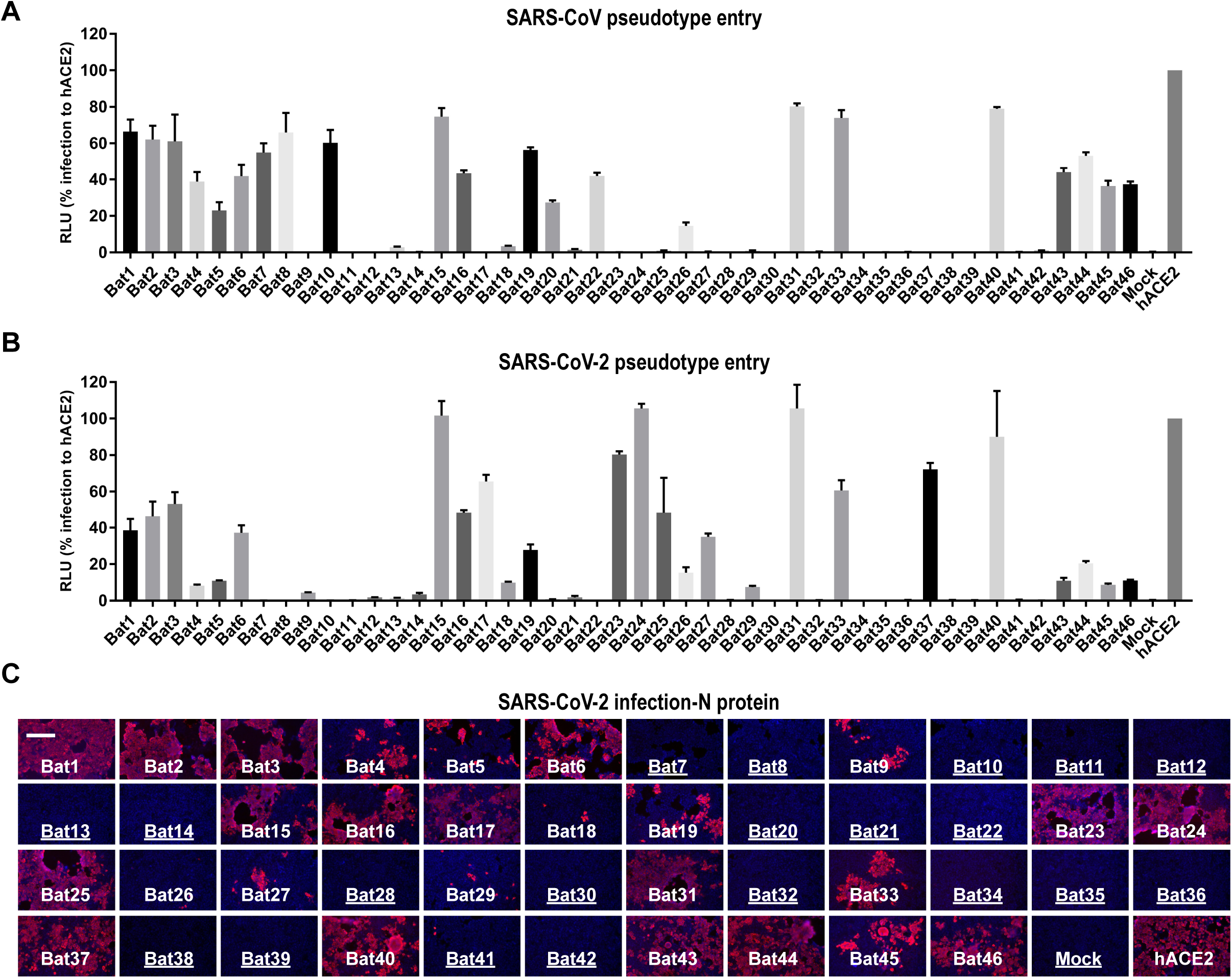
Characterization of bat ACE2 orthologs mediating entry of SARS-CoV and SARS-CoV-2 viruses. (A-B) The ability of bat ACE2 orthologs to support the entry of SARS-CoV and SARS-CoV-2 pseudovirus. 293T cells expressing bat ACE2 orthologs in a 96-well plate were infected with SARS-CoV (A) and SARS-CoV-2 (B) spike protein pseudotyped VSV-dG-Luc. The luciferase activity of the cell lysate was determined at 20 hours post-infection (hpi). (C) 293T cells expressing bat ACE2 orthologs were inoculated with the SARS-CoV-2 live virus at MOI=0.01. N proteins (red) in the infected cells were detected by an immunofluorescence assay at 48 hpi. Scale bar=200 μm. Species that show almost no entry for SARS-CoV-2 live virus are underlined.

The SARS-CoV-2 S protein used here for pseudotyping contains a D614G mutation, which is currently a dominant variation (*16*). The D614G mutation remarkably improved the *in vitro* infectivity of SARS-CoV-2, but may not significantly affect the receptor interaction since it is not in the RBD (*17*). Indeed, we identified a very similar susceptibility profile using an original strain without D614G **(Figure S5)**. We further demonstrated that the pseudotyped entry assay mimics the entry of live viruses through a SARS-CoV-2 infection assay **(Figure 3C)**. As expected, the profile of SARS-CoV-2 N protein expression is highly consistent with the results from the VSV-dG-based pseudotyped virus entry assay **(Figure 3C)**. However, the live virus infection resulted in the phenotype of plaque formation, while the pseudotypes showed evenly distributed single-round infection **(Figure S4)**.

When comparing the RBD-hFC binding and pseudotype entry profiles, we found that binding and susceptibility are generally consistent, with a few exceptions. For instance, some species (Bat12, 13, 14) were able to bind to SARS-CoV-2 RBD-hFc efficiently, but cannot support infection of the same virus, indicating that high binding affinity does not guarantee efficient viral entry **(Figures 2 and 3)**. In contrast, some species (Bat3-8) were defective or less efficient in SARS-CoV RBD-hFc binding, but supported the entry of the same virus to some degree **(Figures 2 and 3)**. We hypothesize that such minimal binding may be sufficient for viral entry mediated by those ACE2 orthologs; alternatively, additional residues outside the traditional RBD region might be required for efficient interaction. These hypotheses should be tested in the future.

Together, our results demonstrated that SARS-CoV and SARS-CoV-2 can selectively use some bat ACE2 as functional receptors for viral entry, but many – if not most – bat ACE2 are not favored by one or both viruses. The functional defects in ACE2 coronavirus receptor in our functional assays provide strong evidence that rejects the suggestion that many/most bat species are potential natural hosts of SARS-CoV and/or SARS-CoV-2.

### Evaluation of critical genetic changes in bat ACE2 orthologs affecting the viral binding and entry efficiency or specificity

We comprehensively analyzed the relationship between critical RBD binding sites in bat ACE2 sequences and their ability to support SARS-CoV and SARS-CoV-2 RBD binding and viral entry. Several critical residues were identified that may play critical roles in the determination of species specificity **(Figure S1)**. According to the sequence alignment, two species pairs (Bat33-34 and Bat38-40) were selected to demonstrate the role of critical residues in RBD binding and viral entry, because they are phylogenetically close but show contrasting phenotypes for supporting RBD binding and viral entry. Specifically, Bat34 and 38 do not support SARS-CoV and SARS-CoV-2 RBD binding and infection, while Bat33 supports efficient binding and infection of both viruses, and Bat40 supports infection of both viruses and to a lesser degree, SARS-RBD binding **(Figures 2 and 3)**. We compared their protein sequences and highlighted the residues that may affect RBD interaction. For example, substitutions I27K, N31G, and K42E were observed when comparing Bat33 and 34, while Q24L, E30K, K35Q, and G354N were present between Bat38 and 40 **(Figure 4A)**. We hypothesized that the discrepancy in binding and infection phenotype is determined by their differences in critical residues for RBD interaction. To test this hypothesis, we designed a residue swap mutagenesis assay to investigate the role of critical residues on RBD binding and virus entry **(Figure 4A)**. We generated four swap mutations and corresponding 293T stable cell lines to test whether these substitutions can achieve the gain-of-function and loss-of-function. All bat ACE2 orthologs and related mutants were expressed at a comparable level after lentiviral transduction, as indicated by the immunofluorescence of the carboxyl-terminal (C-terminal) 3×Flag tag **(Figure 4B)**. Recombinant SARS-CoV and SARS-CoV-2 RBD-hFC proteins were applied to the cells expressing different ACE2, and the binding efficiency was evaluated by fluorescence **(Figure 4C)** and flow cytometry assays **(Figure 4D)**. As expected, the swap of critical residues on the selected four bat ACE2 changed their receptor function to the opposite, except for Bat38m (Bat38 mutant) that remained unable to bind SARS-CoV RBD-hFc **(Figure 4D-4E)**. The GFP **(Figure 4E)** and Luciferase levels **(Figure 4F)** from the pseudotyped virus entry assay, as well as the N protein staining from the live SARS-CoV-2 infection assay **(Figure 4G)** further confirmed our hypothesis at the viral entry level. Structure modeling of bat ACE2 orthologs showed that these residues appeared to occur in the interface between S protein and ACE2 receptor **(Figure 4H-4I)**, and amino acid changes in these sites could potentially lead to different abilities to support RBD binding and viral entry, confirming our results of virus-host receptor binding and infection assays.

**Fig. 4.**
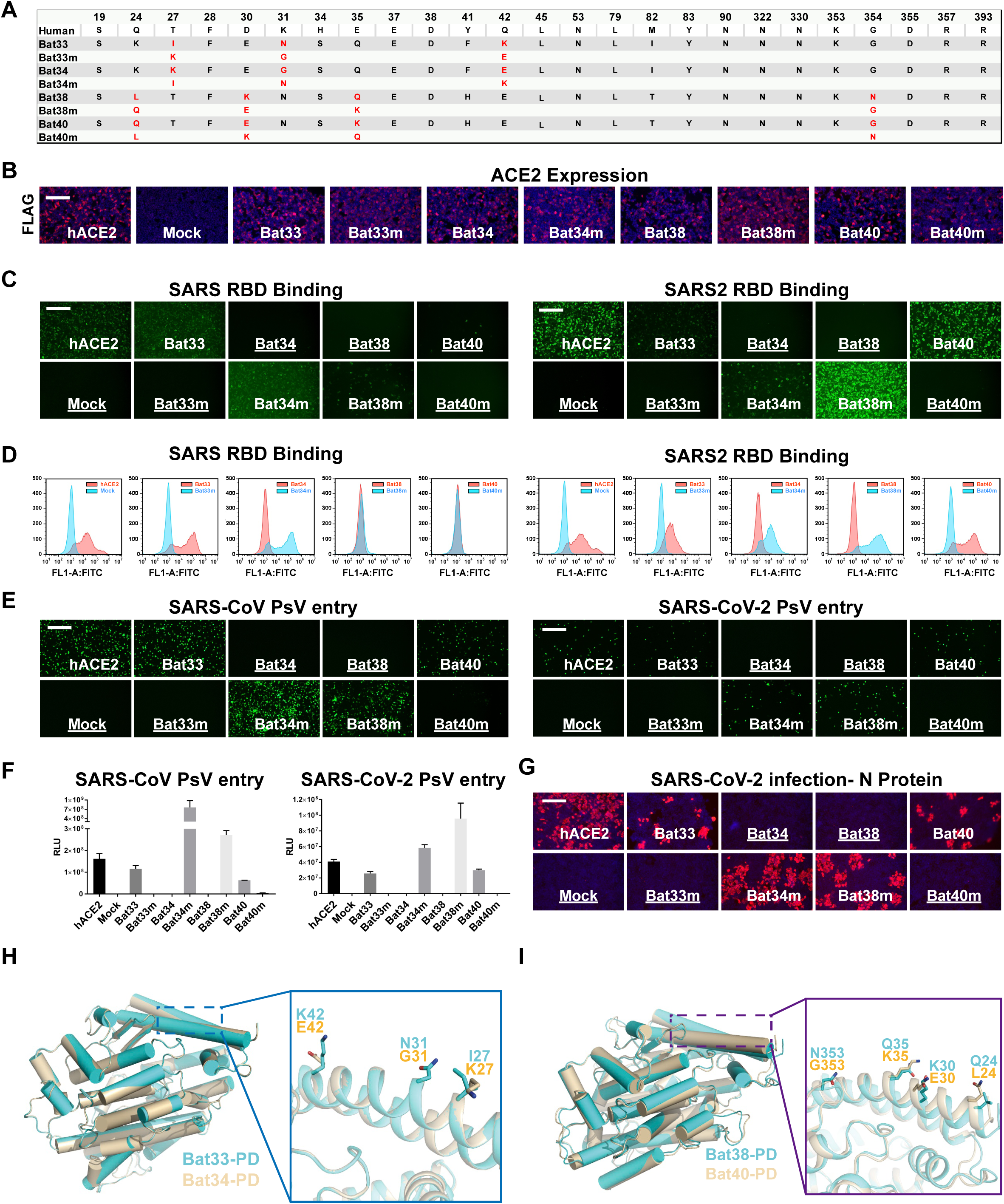
Evaluation of the critical binding sites determining the species-specific restriction of SARS-CoV and SARS-CoV-2 binding and entry. (A) Swap mutagenesis assay to investigate the role of critical binding sites on bat ACE2 orthologs for tropism determination. Residues involved in RBD (according to the structure between SARS2-RBD and human ACE2, PDB: 6M0J) interaction are shown in the table. Residues that changed in the mutagenesis assay are marked in red. (B) The expression level of the bat ACE2 orthologs and related mutants in transduced 293T cells was determined by an immunofluorescence assay recognizing the Flag tag. Scale bar=200 μm. (C-D) Binding efficiency of SARS2-RBD-hFc and SARS2-RBD-hFc on 293T cells expressing bat ACE2 and related mutants. Cells were incubated with 5 μg/ml of recombinant proteins at 37 °C for 1 hour, and then washed and incubated with a secondary antibody recognizing human Fc. Immunostaining (C) and flow cytometry (D) were conducted to show the binding efficiency. Scale bar=200 μm. (E-F) The ability of the indicated ACE2 and related mutants to support the entry of coronavirus pseudotypes. The 293T cells expressing the indicated ACE2 and their mutants were infected by SARS-CoV and SARS-CoV-2 pseudotypes expressing GFP (E) and luciferase (F). Infection was analyzed at 20 hpi. Scale bar=200 μm. (G) 293T cells infected by the SARS-CoV-2 live virus at MOI=0.01; the infection was examined at 48 hpi through N protein (red) immunostaining. Nuclei were stained with Hoechst 33342 in blue. Scale bar=200 μm. (H) Structure alignment for the Bat33 ACE2-PD (cyan) and Bat34 ACE2-PD (wheat). The regions enclosed by the blue-dashed lines are illustrated in detail in the right, in which the variation of the interface residues between Bat33 ACE2-PD (cyan) and Bat34 ACE2-PD (wheat) are indicated by different side chains. (I) Structural alignment for the Bat38 ACE2-PD (cyan) and Bat40 ACE2-PD (wheat). The regions enclosed by the purple dashed lines are illustrated in detail in the right, in which the variation of the interface residues between Bat38 ACE2-PD (cyan) and Bat40 ACE2-PD (wheat) are indicated by different side chains.

## Discussion

Our study provides genetic and functional evidence from bat ACE2 receptor usage to reject the suggestion that many, if not most, bat species are potential hosts of SARS-CoV and SARS-CoV-2. Our sampling covers representative species from 11 bat families, accounting for 96% of all extant bat species, hence providing a broad picture. Moreover, our study included 28 species inhabiting urban areas and 18 species that are not common in cities or do not roost in buildings. Our functional assays demonstrated that there is no correlation between proximity to humans and probability of being natural hosts of SARS-CoV or SARS-CoV-2. Therefore, there is no need to fear the many bat species occurring in cities that are not potential hosts of SARS-CoV and SARS-CoV-2. Species such as horseshoe bats, which are suggested to be potential natural hosts of the two viruses, should also not be feared, as they are less likely to be found in cities.

Our results are only partially consistent with a recently published prediction based on sequence similarity, which estimated a binding score between ACE2 and the SARS-CoV-2 S protein for each vertebrate species examined (*8*). The predicted binding scores for all 37 bat species fell into low (n=8) and very low (n=29) categories (*8*), suggesting that all examined bat species are at low risk for SARS-CoV-2 infection. Our study included 36 of the 37 previously examined bat species **(Figure 1 and Table S1)**; 19 of these appeared to support SARS-CoV-2 entry by their ACE2 receptors **(Figures 1 and 3)**, strongly suggesting that these bats are at high risk for SARS-CoV-2 infection. These disparities between in silico analyses and functional experiments strongly indicate the importance of experimental data for confirmation of in silico analyses, as our understanding of ACE2 sequences and structures is incomplete thus far. Indeed, our genetic and functional evidence revealed critical residues of bat ACE2 that are involved in supporting SARS-CoV-2 entry **(Figure 4)**. However, these residues are not the genetic determinant of New World monkey ACE2 orthologs mediating SARS-CoV-2 entry (*7*), and many bat ACE2 orthologs carrying residues that were considered unfavorable in the same study (H41 and E42) (*7*) were fully functional in our study **(Figure 4)**, further confirming the complexity of ACE2 functionality.

We found that closely related species can show strikingly different ACE2 receptor usage. For example, *Rhinolophus sinicus* can support SARS-CoV entry, whereas its congeneric relatives *R. ferrumequinum* and *R. pearsonii* cannot **(Figures 1 and 3)**, despite the fact that some polymorphic sites of ACE2 may have occurred in *R. sinicus* populations (*18*). These findings clearly show that ACE2 receptor usage is species-dependent. Accordingly, although some bats might be potential hosts of SARS-CoV and SARS-CoV-2 (*1-3*), one cannot assume that all bat species or individuals can carry these viruses. On a positive note, even if some bat species are potential hosts of certain viruses, they do not appear to have overt clinical signs of infection, suggesting that these bats may serve as animal models to develop treatments for humans. Although certain bat species are frequently observed to carry coronaviruses closely related to human viruses in terms of sequence similarity (*19*), there is no solid and direct evidence showing the initial spillover from bats to humans and other animals. Nevertheless, humans infected with coronavirus should maintain distance from bats that can use ACE2 as a viral receptor, because many bat species are endangered and may be susceptible to human coronaviruses (*20*), as suggested for many other mammals (*8, 21*). Indeed, the International Union for Conservation of Nature (IUCN) has assessed that over one third of bat species are threatened or data deficient, and over half of all bat species have unknown or decreasing population trends (*22*). Thus, bats are in need of protection more than ever.

Our study supports the calls that public education on bat biology will reduce the threat to bats (*4, 22*). In fact, all bats are potentially safe as long as they are treated with care and respect. We should work collaboratively to combat the pandemic and identify which species are potential hosts, and not fear those species that are not hosts of the virus. Instead, we must respect and care for those species that are potential hosts, and learn about the impact of human activities on their natural habitats, which may lead to zoonotic spillover events.

## Acknowledgments

We thank B. Fenton, L. Moretoo, and D.M. Morales-Martínez for sharing their knowledge as to whether certain bats roost or forage in cities, Prof. Zheng-Li Shi for providing the SARS-CoV-2 virus, and Ming Dai, Zhixiang Huang, Yan Rao, Jing Zhang, and Bei Wang from ABSL-3 Laboratory of Wuhan University for their technical support. We are grateful to Beijing Taikang Yicai Foundation for their great support to this work.

## Author contributions

H.Z., H.Y., Y.C., and K.L. designed study; H.Z., H.Y., H.J. wrote manuscript; H.Y., H.J., Q.L., Z.Z., X.W., and M.G. performed experiments; H.Y., H.Z., H.J., and Y.C. analyzed data.

## Data and materials availability

All data are available in the manuscript or the supplementary materials.

## Materials and Methods

### ACE2 sequence acquisition and selective pressure analysis

We obtained 46 full-length coding sequences of bat *ACE2* in this study, of which 32 were taken from a recent study (*8*), and 14 were newly extracted from published or recently sequenced genome assemblies (see **Table S1** for the sources and accession numbers for the sequences and assemblies). Next, we aligned the deduced ACE2 protein sequences using the MUSCLE program (*23*) (see **Figure S1** for the resulting alignment). The sequence logo was generated with WebLogo (https://weblogo.berkeley.edu/logo.cgi). We performed selective pressure analysis on bat ACE2 using CodeML implemented in PAML (*24*). Two comparisons of site models (M1a & M2a, M8a & M8) were used to predict positively selected sites (*24*). The input tree was the species tree **(Figure 1)** taken from previous studies (*25-27*).

### Cell culture

HEK293T cells (293T, ATCC, CRL-3216) and VERO-E6 cells (ATCC, CRL-1586) were cultured in Dulbecco’s modified Eagle’s medium (DMEM; Gibco) supplemented with 10% fetal bovine serum (FBS), 2.0 mM L-Glutamine, 110 mg/L sodium pyruvate, and 4.5 g/L D-glucose. l1-Hybridoma (CRL-2700) secreting a monoclonal antibody targeting against VSV glycoprotein was cultured in Minimum Essential Medium with Earle’s salts and 2.0 mM L-Glutamine (MEM; Gibco). All cells were cultured at 37°C in 5% CO_2_ with the regular passage of every 2-3 days. 293T stable cell lines overexpressing ACE2 orthologs were maintained in growth medium supplemented with 1μg/ml puromycin.

### Plasmids

Human codon-optimized cDNA sequences encoding various ACE2 orthologs and their mutants fused with a C-terminus 3×Flag tag (DYKDHD-G-DYKDHD-I-DYKDDDDK) were commercially synthesized and subcloned into a lentiviral transfer vector (pLVX-IRES-puro) through the EcoRI and NotI restriction sites. The DNA sequences of human codon-optimized SARS-CoV S protein (CUHK-W1, GenBank: AY278554.2) and SARS-CoV-2 S protein (Wuhan-Hu-1, GenBank: MN908947) were amplified from plasmids pCMV/hygro-SARS-CoV-S (VG40150-G-N, Sino Biological, China) and pCAGGS-SARS-CoV-2-S-c9 (gifted from Dr. Wenhui Li, National Institute of Biological Science, Beijing, China) into pCAGGS vector with C-terminal 18 aa deletion for improving VSV pseudotyping efficiency (*28, 29*). The D614G mutation was introduced into the SARS-CoV-2-S coding sequence to improve *in vitro* infection efficiency. The plasmids for the expression of coronavirus RBD-IgG Fc fusion proteins were generated by inserting the coding sequences of SARS-CoV RBD (aa 318-516) and SARS-CoV-2 RBD (aa331-530) into the pCAGGS vector to express fusion proteins with C-terminal human Fc (IgG1) and N-terminal CD5 secretion leading sequence (MPMGSLQPLATLYLLGMLVASVL).

### Generation of ACE2 stable expression cell lines

293T cells overexpressing ACE2 orthologs were generated by lentiviral transduction. Specifically, the lentivirus was produced by cotransfection of lentiviral transfer vector carrying ACE2 coding sequences (pLVX-EF1a-Puro, from Genewiz Inc.) and packaging plasmids pMD2G (Addgene #12259) and psPAX2 (Addgene #12260) into 293T cells through Lipofectamine 3000 (Thermo Fisher Scientific, United States). The lentivirus-containing supernatant was collected and pooled at 24 and 48 hours (hrs) post-transfection. HEK293T cells were transduced by the lentivirus after 16 hrs, in the presence of 8 μg/ml polybrene. Stable cells expressing various ACE2 orthologs were selected and maintained in growth medium with puromycin (1 μg/ml).

### Immunofluorescence assay to evaluate the expression level of ACE2 orthologs

The expression levels of ACE2 orthologs were evaluated by the immunofluorescence assay detecting the C-terminal 3×Flag tags. The cells for analysis were seeded in the poly-lysine pretreated 96-well plate at a cell density of 5×10 ^5^/ml (100 μl/well), and cultured for 24 hrs. Cells were fixed with 4% paraformaldehyde at room temperature for 10 mins, permeablized with 0.2% Triton X-100/PBS at room temperature for 10 mins, and blocked with 1% Bovine serum albumin (BSA) at 37°C for 30 mins. Next they were incubated with the mouse monoclonal antibody targeting Flag tag (9A3, #8146S, Cell signaling technology, United States) diluted in 1% BSA/PBS at 37°C for 1 hour. After three rounds of PBS washing, cells were subsequently incubated with 2 μg/ml of the secondary goat anti-rabbit antibody conjugated with Alexa Fluor 594 (A11032, Thermo Fisher Scientific, United States) diluted in 1% BSA /PBS at room temperature for 30 mins. The nucleus was stained with Hoechst 33342 (1:5000 dilution in PBS) in blue. Images were captured with a fluorescence microscope (MI52-N, Mshot, China).

### Production of VSV reporter virus pseudotyped with coronavirus spike proteins

Coronavirus spike protein pseudotyped virus (CoV-psV) were packaged following a previously described protocol using a replicate-deficient VSV based rhabdoviral pseudotyping system (VSV-dG) (*30*). The VSV-G glycoprotein deficient VSV exogenously expressing EGFP (VSV-dG-GFP) or Firefly Luciferase (VSV-dG-Luc) were rescued by a reverse genetics system purchased from a company (Kerafast). To produce CoV-psV, Vero-E6 cells were transfected with the plasmids overexpressing SARS2-CoV (pCAGGS-SARS-S-dc) and SARS2-CoV-2 spike proteins (pCAGGS-SARS2-S-dc) through Lipofectamine 3000 reagent. After 36 hrs, the transfected cells were transduced with VSV-dG reporter viruses diluted in serum-free opti-MEM for 1 hour at 37°C (MOI=10). The transduced cells were washed with culture medium once and then replenished with fresh culture medium with L1 hybridoma cultured supernatant containing anti-VSV mAb (1:100 dilution) to neutralize the infectivity of the residual input viruses. The CoV-psV containing supernatants were harvested at 24 hrs after transduction, clarified at 12,000 rpm for 2 mins at 4°C, and immediately transferred to −80°C for storage. The viral titer (genome equivalents) was determined by quantitative reverse transcription PCR (RT-qPCR). The RNA copies in the virus-containing supernatant were detected using the VSV-L gene sequences.

### Pseudotype entry assay

293T stable cell lines overexpressing various ACE2 orthologs were trypsinized and resuspended together with SARS-CoV or SARS-CoV-2 pseudotyped viruses (at a genome equivalents=100) in DMEM with 10% FBS. Next they were seeded at 5×10 ^4^ in a well of a 96-well plate to allow attachment and viral infection simultaneously. At 16-24 hrs after infection, images of infected cells with GFP expression were acquired with a fluorescence microscope (MI52-N, Mshot, China). Cells infected with pseudovirus expressing firefly luciferase were lyzed by 1× passive lysis buffer (Promega, United States) at room temperature for 15 mins. Luciferase activity in the cell lysate was determined by a Bright-Glo luciferase assay kit (Promega, United States) and measured through a Spectra MaxiD3 multi-well Luminometer (Molecular Devices, United States) or a GloMax® 20/20 Luminometer (Promega, United States).

### Coronavirus RBD-hFc binding assay

Recombinant SARS-CoV-RBD-hFc and SARS-CoV-2-RBD-hFc proteins were produced by transient transfection of 293T cells with Lipofectamine 3000. The transfected cells were cultured in Free-style 293 serum-free medium (Thermo Scientific), and the supernatants containing the recombinant proteins were collected at 2 and 4 days post-transfection. The RBD-hFc protein concentration was determined by comparing the target protein band with BSA standard dilutions through Coomassie staining. The RBD-hFc protein-containing supernatant was diluted with culture medium (5-10 μg/ml) and then incubated with the 293T stable cell line overexpressing different ACE2 orthologs for 1 hour at 37°C. Cells were washed twice with DMEM and then incubated with 2 μg/ml of Alexa Fluor 488 conjugated Goat anti-Human IgG (A11013, Thermo Fisher Scientific, United States) diluted in DMEM with 2% FBS for 30 mins at 37°C. For immunostaining, cells were washed twice with PBS and incubated with PBS with Hoechst 33342 (1:5000 dilution in PBS) for nucleus staining. Images were captured with a fluorescence microscope (MI52-N, Mshot, China). For flow cytometry analysis, cells were detached by 5mM EDTA/PBS and analyzed with a CytoFLEX Flow Cytometer (Beckman Coulter, United States).

### SARS-CoV-2 live virus infection assay

The SARS-CoV-2 (strain IVCAS 6.7512) was provided by the National Virus Resource, Wuhan Institute of Virology, Chinese Academy of Sciences. All SARS-CoV-2 live virus related experiments were approved by the Biosafety Committee Level 3 (ABSL-3) of Wuhan University. All experiments involving SARS-CoV-2 were performed in the BSL-3 facility. SARS-CoV-2 was amplified on Vero-E6 cells and stored at −150°C, and the titer was determined on Vero-E6 cells through a plaque assay. 293T cells expressing ACE2 orthologs were seeded on a poly-lysine coated 96-well plate for 24 hrs before inoculation. Cells were infected with SARS-CoV-2 at MOI=0.01, and then incubated in DMEM with 2% FBS for 48 hrs before testing. The cells were fixed with 4% paraformaldehyde in PBS at room temperature for 1 hour, permeablized with 0.2% Triton X-100 for 10 mins, and then blocked with 1% BSA/PBS at 37°C for 1 hour. Cells were subsequently incubated with a mouse monoclonal antibody SARS-CoV/SARS-CoV-2 Nucleocapsid Antibody (40143-MM05, Sino Biological, China) at 1:500 at 37°C for 1 hour, and then incubated with 2μg/ml of goat anti-mouse secondary antibody, Alexa Fluor 594 (A-11032, Thermo Fisher Scientific) at 37°C for 1 hour. The nucleus was stained with Hoechst 33342. Images were acquired with a fluorescence microscope (MI52-N, Mshot, China).

### Homology-based structural modeling

Molecular models of different bat ACE2 were predicted by I-TASSER (Iterative Threading ASSEmbly Refinement) version 5.1 (*31*). Starting from the amino acid sequences, the I-TASSER algorithm constructed the full-length 3D atomic models by structural template identification, followed by template-based fragment assembly simulations. The model with the highest confidence score in each prediction was used for subsequent analyses (*32*). Only the predicted structures of the N-terminal peptidase domain (PD) of ACE2 were used in the analyses. The structural alignment and visualization were implemented in PyMOL (*33*).

### Statistical analysis

Data are expressed as mean values with standard deviation. All experiments were repeated 3-5 times, each yielding similar results.

