## Supplementary Tables and Figures for "Many bat species are not potential hosts of SARS-CoV and SARS-CoV-2: Evidence from ACE2 receptor usage"

**Affiliations:**

**This PDF file includes:**

Supplementary Tables S1-S5

Supplementary Figures S1-S5

**Table S1. Data source of bat ACE2 sequences.**

| Family | Species | Label | Source | Reference |
|---|---|---|---|---|
| Pteropodidae | *Rousettus aegyptiacus* | Bat01 | XM_016118926.1 (NCBI gene) | Damas et al. 2020 PNAS |
| | *Pteropus alecto* | Bat02 | XM_006911647.1 (NCBI gene) | Damas et al. 2020 PNAS |
| | ***Pteropus giganteus*** | **Bat03** | **GCA_902729225.1 (NCBI assembly)** | **This study** |
| | *Eidolon helvum* | Bat04 | GCA_000465285.1 (NCBI assembly) | Damas et al. 2020 PNAS |
| | *Eonycteris spelaea* | Bat05 | GCA_003508835.1 (NCBI assembly) | Damas et al. 2020 PNAS |
| | *Macroglossus sobrinus* | Bat06 | GCA_004027375.1 (NCBI assembly) | Damas et al. 2020 PNAS |
| | *Cynopterus sphinx* | Bat07 | MT515623 (NCBI gene) | Damas et al. 2020 PNAS |
| | ***Cynopterus brachyotis*** | **Bat08** | **GCA_009793145.1 (NCBI assembly)** | **This study** |
| Rhinolophidae | *Rhinolophus ferrumequinum* | Bat09 | GCA_004115265.2 (NCBI assembly) | Damas et al. 2020 PNAS |
| | *Rhinolophus sinicus* | Bat10 | XM_019746337.1 (NCBI gene) | Damas et al. 2020 PNAS |
| | *Rhinolophus pearsonii* | Bat11 | MT515622 (NCBI gene) | Damas et al. 2020 PNAS |
| Hipposideridae | *Hipposideros armiger* | Bat12 | XM_019667391.1 (NCBI gene) | Damas et al. 2020 PNAS |
| | *Hipposideros galeritus* | Bat13 | GCA_004027415.1 (NCBI assembly) | Damas et al. 2020 PNAS |
| | *Hipposideros pratti* | Bat14 | MT515621 (NCBI gene) | Damas et al. 2020 PNAS |
| Megadermatidae | *Megaderma lyra* | Bat15 | MT515624 (NCBI gene) | Damas et al. 2020 PNAS |
| Noctilionidae | *Noctilio leporinus* | Bat16 | GCA_004026585.1 (NCBI assembly) | Damas et al. 2020 PNAS |
| Emballonuridae | ***Taphozous melanopogon*** | **Bat17** | **MT952961 (NCBI gene)** | **This study** |
| Phyllostomidae | *Anoura caudifer* | Bat18 | GCA_004027475.1 (NCBI assembly) | Damas et al. 2020 PNAS |
| | ***Trachops cirrhosus*** | **Bat19** | **MT952962 (NCBI gene)** | **This study** |
| | ***Vampyram spectrum*** | **Bat20** | **MT952963 (NCBI gene)** | **This study** |
| | *Tonatia saurophila* | Bat21 | GCA_004024845.1 (NCBI assembly) | Damas et al. 2020 PNAS |
| | *Phyllostomus discolor* | Bat22 | XM_028522516.1 (NCBI gene) | Damas et al. 2020 PNAS |
| | ***Carollia perspicillata*** | **Bat23** | **GCA_004027735.1 (NCBI assembly)** | **This study** |
| | *Micronycteris hirsuta* | Bat24 | GCA_004026765.1 (NCBI assembly) | Damas et al. 2020 PNAS |
| | ***Sturnira hondurensis*** | **Bat25** | **GWHAAZA00000000 (Genome Warehouse assembly)** | **This study** |
| | ***Artibeus jamaicensis*** | **Bat26** | **GCA_004027435.1 (NCBI assembly)** | **This study** |
| | *Desmodus rotundus* | Bat27 | XM_024569930.1 (NCBI gene) | Damas et al. 2020 PNAS |

| | | | | |
|---|---|---|---|---|
| Mormoopidae | *Pteronotus parnellii* | Bat28 | GCA_000465405.1 (NCBI assembly) | Damas et al. 2020 PNAS |
| | *Mormoops blainvillei* | Bat29 | GCA_004026545.1 (NCBI assembly) | Damas et al. 2020 PNAS |
| | ***Pteronotus davyi*** | **Bat30** | **MT952964 (NCBI gene)** | **This study** |
| Molossidae | *Tadarida brasiliensis* | Bat31 | GCA_004025005.1 (NCBI assembly) | Damas et al. 2020 PNAS |
| | *Molossus molossus* | Bat32 | https://vgp.github.io/genomeark/Molossus_molossus (genome) | Damas et al. 2020 PNAS |
| Miniopteridae | *Miniopterus schreibersii* | Bat33 | GCA_004026525.1 (NCBI assembly) | Damas et al. 2020 PNAS |
| | *Miniopterus natalensis* | Bat34 | GCA_001595765.1 (NCBI assembly) | Damas et al. 2020 PNAS |
| Vespertilionidae | *Eptesicus fuscus* | Bat35 | XM_008154928.2 (NCBI gene) | Damas et al. 2020 PNAS |
| | ***Aeorestes*** **(*Lasiurus*)** ***cinereus*** | **Bat36** | **GCA_011751065.1 (NCBI assembly)** | **This study** |
| | ***Pipistrellus pipistrellus*** | **Bat37** | **GCA_004026625.1 (NCBI assembly)** | **This study** |
| | ***Lasiurus borealis*** | **Bat38** | **GCA_004026805.1 (NCBI assembly)** | **This study** |
| | *Pipistrellus kuhlii* | Bat39 | https://vgp.github.io/genomeark/Pipistrellus_kuhlii (genome) | Damas et al. 2020 PNAS |
| | ***Antrozous pallidus*** | **Bat40** | **GCA_007922775.1 (NCBI assembly)** | **This study** |
| | ***Nycticeius humeralis*** | **Bat41** | **GCA_007922795.1 (NCBI assembly)** | **This study** |
| | *Murina feae* | Bat42 | GCA_004026665.1 (NCBI assembly) | Damas et al. 2020 PNAS |
| | *Myotis myotis* | Bat43 | https://vgp.github.io/genomeark/Myotis_myotis (genome) | Damas et al. 2020 PNAS |
| | *Myotis davidii* | Bat44 | XM_006775210.2 (NCBI gene) | Damas et al. 2020 PNAS |
| | *Myotis brandtii* | Bat45 | XM_014544294.1 (NCBI gene) | Damas et al. 2020 PNAS |
| | *Myotis lucifugus* | Bat46 | XM_023753669.1 (NCBI gene) | Damas et al. 2020 PNAS |

**Table S2. Roosting and foraging sites of the 46 bat species.**

| Family | Species | Label | Roosting site | Foraging site | In close contact with humans? |
|---|---|---|---|---|---|
| Pteropodidae | *Rousettus aegyptiacus* | Bat01 | urban caves | urban area | yes |
| | *Pteropus alecto* | Bat02 | urban forests | urban area | yes |
| | *Pteropus giganteus* | Bat03 | urban forests | urban area | yes |
| | *Eidolon helvum* | Bat04 | urban forests | urban area | yes |
| | *Eonycteris spelaea* | Bat05 | urban caves | urban area | yes |
| | *Macroglossus sobrinus* | Bat06 | urban forests | urban area | yes |
| | *Cynopterus sphinx* | Bat07 | urban trees | urban area | yes |
| | *Cynopterus brachyotis* | Bat08 | urban trees | urban area | yes |
| Rhinolophidae | *Rhinolophus ferrumequinum* | Bat09 | rural caves, buildings | rural area | no |
| | *Rhinolophus sinicus* | Bat10 | rural caves | rural area | no |
| | *Rhinolophus pearsonii* | Bat11 | rural caves | rural area | no |
| Hipposideridae | *Hipposideros armiger* | Bat12 | rural caves | rural area | no |
| | *Hipposideros galeritus* | Bat13 | rural caves | rural area | no |
| | *Hipposideros pratti* | Bat14 | rural caves | rural area | no |
| Megadermatidae | *Megaderma lyra* | Bat15 | rural caves | rural area | no |
| Noctilionidae | *Noctilio leporinus* | Bat16 | urban caves | rural area | yes |
| Emballonuridae | *Taphozous melanopogon* | Bat17 | urban caves | rural area | yes |
| Phyllostomidae | *Anoura caudifer* | Bat18 | rural forests | rural area | no |
| | *Trachops cirrhosus* | Bat19 | rural forests | rural area | no |
| | *Vampyram spectrum* | Bat20 | rural forests | rural area | no |
| | *Tonatia saurophila* | Bat21 | rural forests | rural area | no |
| | *Phyllostomus discolor* | Bat22 | urban caves | urban area | yes |
| | *Carollia perspicillata* | Bat23 | urban forests, caves | urban area | yes |
| | *Micronycteris hirsuta* | Bat24 | rural forests | rural area | no |
| | *Sturnira hondurensis* | Bat25 | rural forests | rural area | no |
| | *Artibeus jamaicensis* | Bat26 | urban forests | urban area | yes |

| | | | | | |
|---|---|---|---|---|---|
| | *Desmodus rotundus* | Bat27 | urban caves | urban area | yes |
| Mormoopidae | *Pteronotus parnellii* | Bat28 | rural caves | rural area | no |
| | *Mormoops blainvillei* | Bat29 | rural caves | rural area | no |
| | *Pteronotus davyi* | Bat30 | rural caves | rural area | no |
| Molossidae | *Tadarida brasiliensis* | Bat31 | urban caves, crevices | urban area | yes |
| | *Molossus molossus* | Bat32 | urban caves, roofs | urban area | yes |
| Miniopteridae | *Miniopterus schreibersii* | Bat33 | rural caves, thatches | rural area | no |
| | *Miniopterus natalensis* | Bat34 | rural caves, thatches | rural area | no |
| Vespertilionidae | *Eptesicus fuscus* | Bat35 | urban caves, buildings | urban area | yes |
| | *Aeorestes* (*Lasiurus*) *cinereus* | Bat36 | urban foliage | urban area | yes |
| | *Pipistrellus pipistrellus* | Bat37 | urban caves, buildings | urban area | yes |
| | *Lasiurus borealis* | Bat38 | urban foliage | urban area | yes |
| | *Pipistrellus kuhlii* | Bat39 | urban caves | urban area | yes |
| | *Antrozous pallidus* | Bat40 | urban caves | urban area | yes |
| | *Nycticeius humeralis* | Bat41 | urban caves, buildings | urban area | yes |
| | *Murina feae* | Bat42 | urban caves | urban area | yes |
| | *Myotis myotis* | Bat43 | urban caves, buildings | urban area | yes |
| | *Myotis davidii* | Bat44 | urban caves | urban area | yes |
| | *Myotis brandtii* | Bat45 | urban caves | urban area | yes |
| | *Myotis lucifugus* | Bat46 | urban caves, buildings | urban area | yes |

Note- "yes" indicates that these species were found to coexist with humans, inclusive in large cities, suggesting that these bats are in close contact with humans. "no" indicates that these species probably do not occur in cities or roost in human buildings and are more restricted to forests, compared to most vespertilionid species that occur in cities more frequently, suggesting that these bats are probably in distant contact with humans.

**Table S3. Species numbers from each bat family.** Data are taken from Wilson and Mittermeier (2019). Bat families that contain species examined in this study are shown in bold.

| | Family | Species number |
|---|---|---|
| 1 | **Pteropodidae** | 191 |
| 2 | Rhinopomatidae | 6 |
| 3 | Craseonycteridae | 1 |
| 4 | **Megadermatidae** | 6 |
| 5 | Rhinonycteridae | 9 |
| 6 | **Hipposideridae** | 88 |
| 7 | **Rhinolophidae** | 109 |
| 8 | **Emballonuridae** | 54 |
| 9 | Nycteridae | 15 |
| 10 | Myzopodidae | 2 |
| 11 | Mystacinidae | 2 |
| 12 | **Noctilionidae** | 2 |
| 13 | Furipteridae | 2 |
| 14 | Thyropteridae | 5 |
| 15 | **Mormoopidae** | 18 |
| 16 | **Phyllostomidae** | 217 |
| 17 | Natalidae | 12 |
| 18 | **Molossidae** | 126 |
| 19 | **Miniopteridae** | 38 |
| 20 | Cistugidae | 2 |
| 21 | **Vespertilionidae** | 496 |
| | Subtotal | 1401 |

Note- Bats have a total of 21 families that contain 1401 species. In this study, we examined 46 species representing 11 families that contain 1345 species, accounting for 96% of all bat species; whereas the remaining 10 families contain only 56 species, accounting for 4% of all bat species.

**Table S4**. **Prediction of positively selected sites for bat *ACE2*.**

| **Site models** | **np**[a] | **Ln *L***[b] | **Model compared** | ***P* value**[c] | **Positively selected sites**[d] |
|---|---|---|---|---|---|
| **M1a** | 93 | -22716.416 | | | |
| **M2a** | 95 | -22540.387 | M1a vs M2a | 3.56E-77 | **24L** (0.999), **27T** (1.000), **82T** (1.000), 91P (1.000), 92E (0.998), 212I (1.000), 213N (1.000), 299N (0.968), **354G** (0.999), 387T (1.000), 429Y (1.000), 531R (0.976), 552K (0.974), 568L (1.000), 572S (1.000), 607S (1.000), 656L (0.977), 658V (1.000), 671W (1.000), 675L (1.000), 689Q (1.000), 691V (1.000) |
| **M8a** | 94 | -22688.470 | | | |
| **M8** | 95 | -22540.956 | M8a vs M8 | 3.99E-66 | **24L** (1.000), **27T** (1.000), **82T** (1.000), 91P (1.000), 92E (0.999), 212I (1.000), 213N (1.000), 299N (0.985), 316V (0.967), 321P (0.968), **354G** (1.000), 387T (1.000), 429Y (1.000), 531R (0.991), 552K (0.988), 568L (1.000), 572S (1.000), 607S (1.000), 644 K (0.978), 656L (0.992), 658V (1.000), 671W (1.000), 675L (1.000), 689Q (1.000), 691V (1.000) |

[a] Numbers of parameters.

[b] The natural logarithm of the likelihood value.

[c] *P* values were generated by comparing the two models with a chi-square test.

[d] Positively selected sites with the posterior probabilities >0.95 were listed based on bayes empirical bayes (BEB) analysis. Sites shown in bold were identified as key residues involved in binding to SARS-CoV-2 S protein.

**Table S5. Infection abilities of bat ACE2 receptors to support SARS-CoV and SARS-CoV-2 entry.** Infection data were indicated as % mean values of bat ACE2 supporting infection compared with the infection supported by human ACE2. The infection efficiency smaller than 5% is shown in red, between 5% and 50% in black, and greater than 5% in green.

| Species | Label | SARS-CoV | SARS-CoV-2 |
|---|---|---|---|
| ***Rousettus aegyptiacus*** | Bat01 | **68.89%** | **38.53%** |
| ***Pteropus alecto*** | Bat02 | **63.56%** | 46.46% |
| ***Pteropus giganteus*** | Bat03 | **64.82%** | **53.28%** |
| ***Eidolon helvum*** | Bat04 | 40.20% | 8.18% |
| ***Eonycteris spelaea*** | Bat05 | 24.16% | 10.92% |
| ***Macroglossus sobrinus*** | Bat06 | 42.79% | 37.26% |
| ***Cynopterus sphinx*** | Bat07 | **55.67%** | **0.12%** |
| ***Cynopterus brachyotis*** | Bat08 | **70.35%** | **0.07%** |
| ***Rhinolophus ferrumequinum*** | Bat09 | **0.23%** | **4.43%** |
| ***Rhinolophus sinicus*** | Bat10 | **60.12%** | **0.09%** |
| ***Rhinolophus pearsonii*** | Bat11 | **0.14%** | **0.21%** |
| ***Hipposideros armiger*** | Bat12 | **0.01%** | **1.88%** |
| ***Hipposideros galeritus*** | Bat13 | **2.76%** | **1.59%** |
| ***Hipposideros pratti*** | Bat14 | **0.31%** | **3.49%** |
| ***Megaderma lyra*** | Bat15 | **72.19%** | **101.65%** |
| ***Noctilio leporinus*** | Bat16 | 42.72% | 48.48% |
| ***Taphozous melanopogon*** | Bat17 | **0.16%** | **65.58%** |
| ***Anoura caudifer*** | Bat18 | **3.39%** | 9.82% |
| ***Trachops cirrhosus*** | Bat19 | **55.63%** | 27.76% |
| ***Vampyram spectrum*** | Bat20 | 26.88% | **0.75%** |
| ***Tonatia saurophila*** | Bat21 | **1.55%** | **1.83%** |
| ***Phyllostomus discolor*** | Bat22 | 41.03% | **0.17%** |
| ***Carollia perspicillata*** | Bat23 | **0.11%** | **80.32%** |
| ***Micronycteris hirsuta*** | Bat24 | **0.02%** | **105.57%** |
| ***Sturnira hondurensis*** | Bat25 | **0.99%** | 48.42% |
| ***Artibeus jamaicensis*** | Bat26 | 15.45% | 15.38% |
| ***Desmodus rotundus*** | Bat27 | **0.48%** | 35.07% |
| ***Pteronotus parnellii*** | Bat28 | **0.27%** | **0.38%** |
| ***Mormoops blainvillei*** | Bat29 | **1.03%** | 7.44% |
| ***Pteronotus davyi*** | Bat30 | **0.17%** | **0.25%** |
| ***Tadarida brasiliensis*** | Bat31 | **80.04%** | 105.51% |
| ***Molossus molossus*** | Bat32 | **0.49%** | **0.32%** |
| ***Miniopterus schreibersii*** | Bat33 | **72.62%** | **60.61%** |
| ***Miniopterus natalensis*** | Bat34 | **0.27%** | **0.14%** |
| ***Eptesicus fuscus*** | Bat35 | **0.09%** | **0.24%** |
| ***Aeorestes (Lasiurus) cinereus*** | Bat36 | **0.33%** | **0.39%** |
| ***Pipistrellus pipistrellus*** | Bat37 | **0.02%** | **72.23%** |
| ***Lasiurus borealis*** | Bat38 | **0.05%** | **0.31%** |
| ***Pipistrellus kuhlii*** | Bat39 | **0.16%** | **0.31%** |

| | | | |
|---|---|---|---|
| ***Antrozous pallidus*** | Bat40 | **78.63%** | **90.09%** |
| ***Nycticeius humeralis*** | Bat41 | **0.29%** | **0.57%** |
| ***Murina feae*** | Bat42 | **1.05%** | **0.09%** |
| ***Myotis myotis*** | Bat43 | 44.57% | 10.91% |
| ***Myotis davidii*** | Bat44 | **52.08%** | 20.46% |
| ***Myotis brandtii*** | Bat45 | 35.08% | 8.70% |
| ***Myotis lucifugus*** | Bat46 | 36.54% | 11.07% |
| **Human** | hACE2 | 100.00% | 100.00% |
| **Mock** | Mock | 0.35% | 0.29% |

**Legends of supplemental figures:**

**Figure S1. Sequence alignment of RBD interacting region on 46 bat ACE2 orthologs.** The alignment of RBD interaction regions were generated by MEGA-X software. Amino acids were colored based on their residue features. The residues involved in the interaction between human ACE2 and SARS-CoV-2 RBD were indicated on the top. Sequence logo of the corresponding regions was generated by WebLogo (bottom).

**Figure S2. Flow cytometry analysis of the SARS-CoV-2 RBD-hFc binding.** 293T cells expressing bat ACE2 orthologs were incubated with 5 μg/ml of recombinant SARS-CoV-2 RBD-hFc protein at 37℃ for 1 hour, and then washed and incubated with a Alexa Fluor 488 conjugated secondary antibody recognizing human IgG Fc. Histogram charts were generated by FlowJo with mock cells (sample 47) as negative control.

**Figure S3. Verification of SARS-CoV and SARS-CoV-2 pseudotypes on 293T cells expressing human ACE2.** A high signal/background ratio of viral entry can be achieved on 293T-hACE2 cells for SARS-CoV and SARS-CoV-2 spike protein pseudotyped VSV-dG viruses expressing Firefly Luciferase (left) and GFP (right).

**Figure S4. The ability of bat ACE2 orthologs to support the entry of SARS-CoV and SARS-CoV-2 pseudotyped VSV-dG-GFP viruses.** 293T cells expressing the ACE2 orthologs of the indicated bats were infected with SARS-CoV and SARS-CoV-2 pseudotyped VSV-dG-GFP viruses. Images were captured at 20 hpi. Scale bar=200 μm.

**Figure S5. Infection profile of virus pseudotyped with SARS-CoV-2 S protein without D614G variation.** 293T cells expressing the indicated bats ACE2 orthologs were infected with SARS-CoV-2 (Wuhan-Hu-1 strain, without D614G variation) S protein pseudotyped VSV-dG-luc viruses. Luciferase units were determined at 20 hpi. Scale bar=200 μm.

### Figure S1

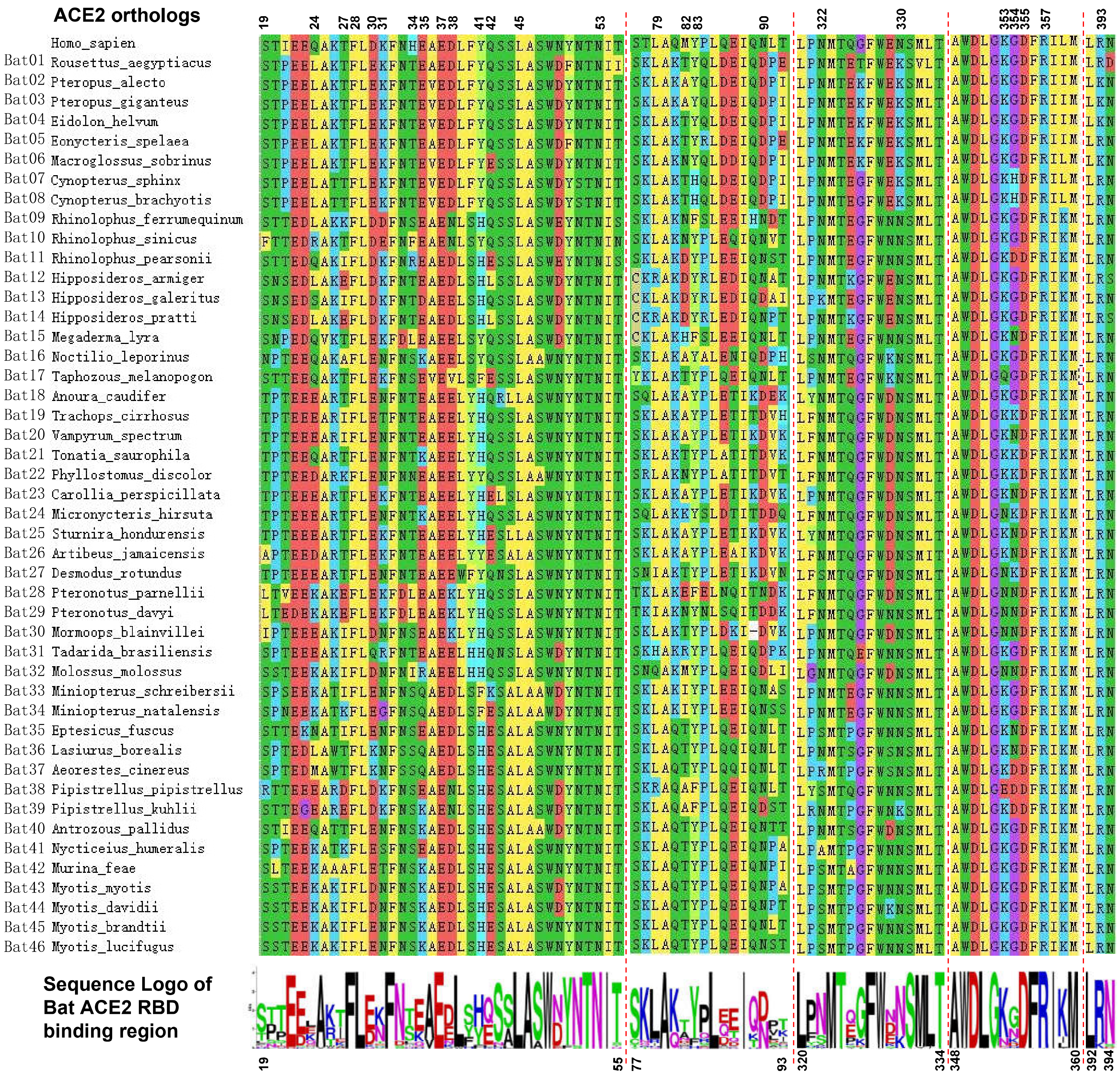
ACE2 orthologs
19 24 27 28 30 31 34 35 37 38 41 42 45 53 79 82 83 90 322 330 353 354 355 357 393
Homo_sapien
Bat01 Rousettus_aegyptiacus
Bat02 Pteropus_alecto
Bat03 Pteropus_giganteus
Bat04 Eidolon_helvum
Bat05 Eonycteris_spelaea
Bat06 Macroglossus_sobrinus
Bat07 Cynopterus_sphinx
Bat08 Cynopterus_brachyotis
Bat09 Rhinolophus_ferrumequinum
Bat10 Rhinolophus_sinicus
Bat11 Rhinolophus_pearsonii
Bat12 Hipposideros_armiger
Bat13 Hipposideros_galeritus
Bat14 Hipposideros_pratti
Bat15 Megaderma_lyra
Bat16 Noctilio_leporinus
Bat17 Taphozous_melanopogon
Bat18 Anoura_caudifer
Bat19 Trachops_cirrhosus
Bat20 Vampyrum_spectrum
Bat21 Tonatia_saurophila
Bat22 Phyllostomus_discolor
Bat23 Carollia_perspicillata
Bat24 Micronycteris_hirsuta
Bat25 Sturnira_hondurensis
Bat26 Artibeus_jamaicensis
Bat27 Desmodus_rotundus
Bat28 Pteronotus_parnellii
Bat29 Pteronotus_davyi
Bat30 Mormoops_blainvillei
Bat31 Tadarida_brasiliensis
Bat32 Molossus_molossus
Bat33 Miniopterus_schreibersii
Bat34 Miniopterus_natalensis
Bat35 Eptesicus_fuscus
Bat36 Lasiurus_borealis
Bat37 Aeorestes_cinereus
Bat38 Pipistrellus_pipistrellus
Bat39 Pipistrellus_kuhlii
Bat40 Antrozous_pallidus
Bat41 Nycticeius_humeralis
Bat42 Murina_feae
Bat43 Myotis_myotis
Bat44 Myotis_davidii
Bat45 Myotis_brandtii
Bat46 Myotis_lucifugus
Sequence Logo of Bat ACE2 RBD binding region
19 55 77 93 320 334 348 360 392 394

### Figure S2

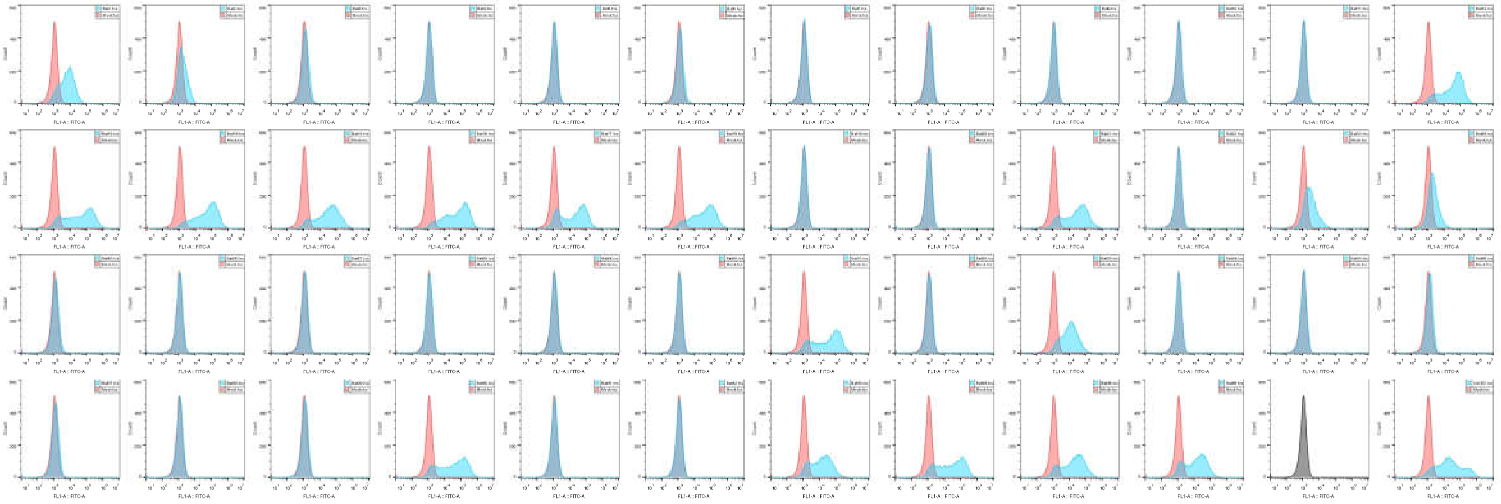

### Figure S3

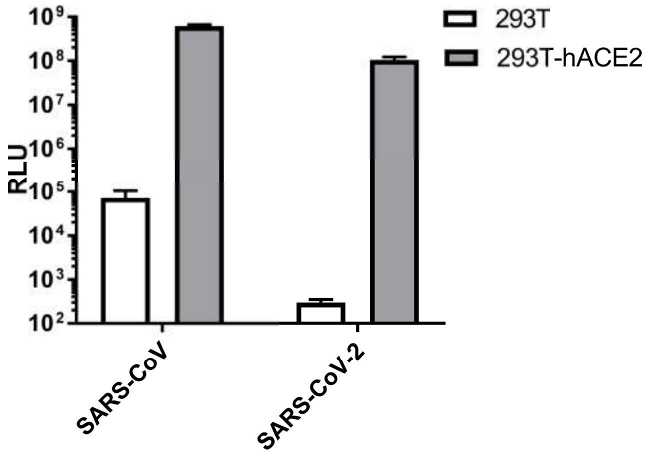

293T
293T-hACE2
RLU
$10^9$
$10^8$
$10^7$
$10^6$
$10^5$
$10^4$
$10^3$
$10^2$
SARS-CoV
SARS-CoV-2


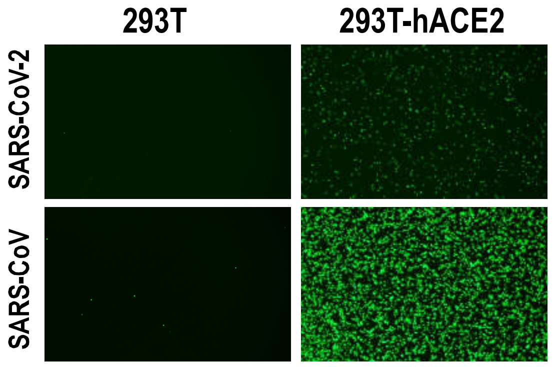

293T
293T-hACE2
SARS-CoV-2
SARS-CoV

### Figure S4

SARS-CoV-GFP infection

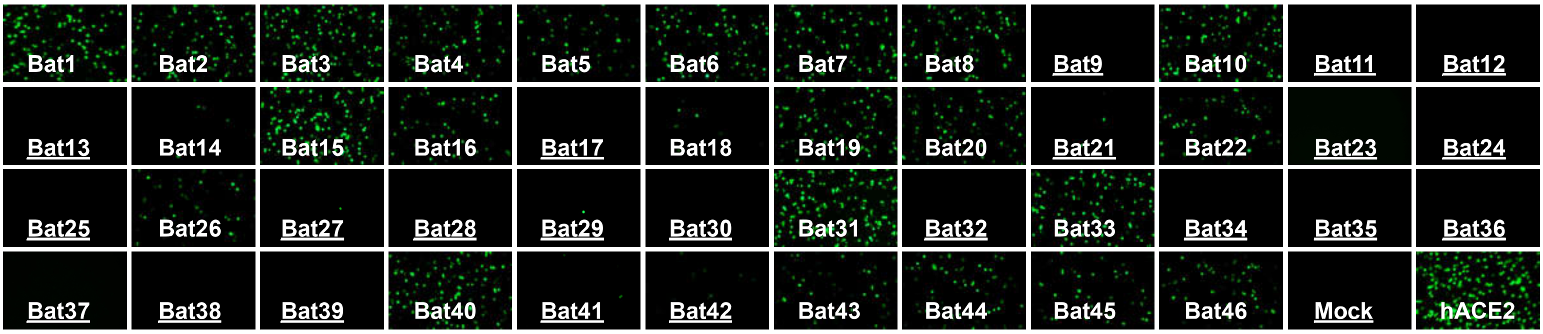


SARS-CoV-2-GFP infection

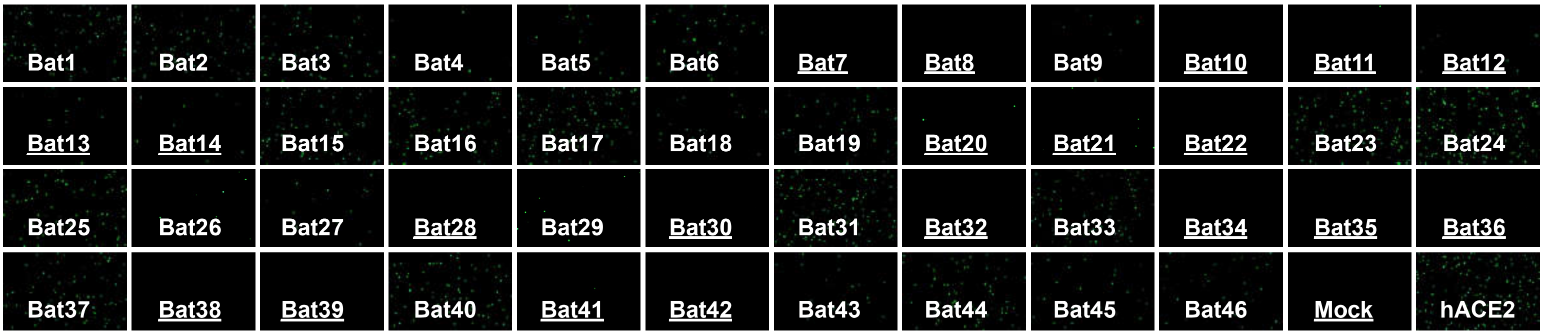

### Figure S5

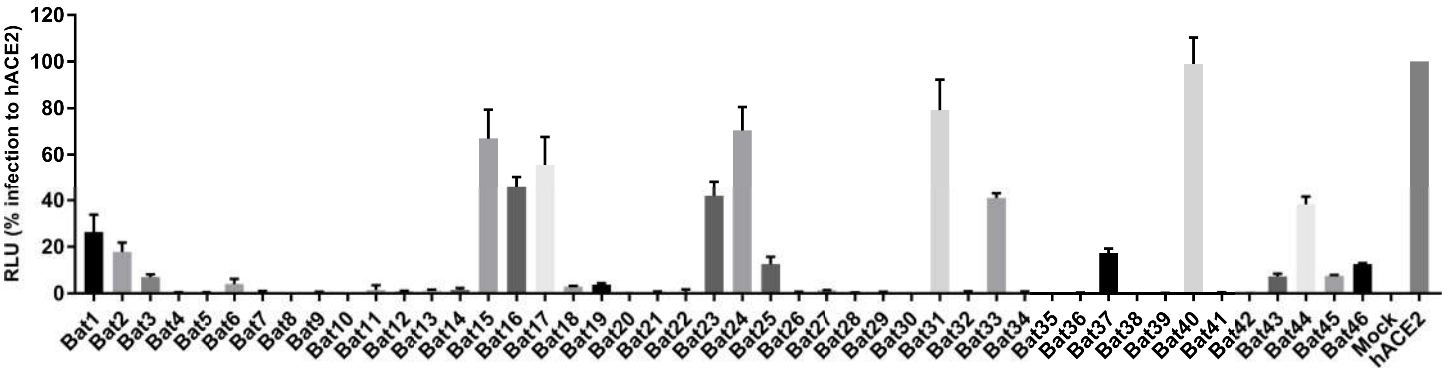

RLU (% infection to hACE2)
120
100
80
60
40
20
0
Bat1
Bat2
Bat3
Bat4
Bat5
Bat6
Bat7
Bat8
Bat9
Bat10
Bat11
Bat12
Bat13
Bat14
Bat15
Bat16
Bat17
Bat18
Bat19
Bat20
Bat21
Bat22
Bat23
Bat24
Bat25
Bat26
Bat27
Bat28
Bat29
Bat30
Bat31
Bat32
Bat33
Bat34
Bat35
Bat36
Bat37
Bat38
Bat39
Bat40
Bat41
Bat42
Bat43
Bat44
Bat45
Bat46
Mock
hACE2
